# A neuromechanical model accounting for movement history dependency identifies subject-specific neural and non-neural origins of joint hyper-resistance: A simulation study

**DOI:** 10.1101/2023.11.09.566428

**Authors:** Jente Willaert, Kaat Desloovere, Anja Van Campenhout, Lena H. Ting, Friedl De Groote

**Affiliations:** KU Leuven; KU Leuven / UZ Leuven; Emory University and Georgia Tech

## Abstract

Joint hyper-resistance is a common symptom in neurological disorders. It has both neural and nonneural origins, but it has been challenging to distinguish different origins based on clinical tests alone. Combining instrumented tests with parameter identification based on a neuromechanical model may allow us to dissociate the different origins of joint hyper-resistance in individual patients. However, this requires that the model captures the underlying mechanisms. Here, we propose a neuromechanical model that, in contrast to previously proposed models, accounts for muscle shortrange stiffness and its interaction with muscle tone and reflex activity.

We collected knee angle trajectories during the pendulum test in 15 children with cerebral palsy (CP) and 5 typically developing children. We did the test in two conditions – hold and pre-movement – that have been shown to alter knee movement. We modeled the lower leg as an inverted pendulum actuated by two antagonistic Hill-type muscles extended with SRS. Reflex activity was modeled as delayed, linear feedback from muscle force. We estimated neural and non-neural parameters by optimizing the fit between simulated and measured knee angle trajectories during the hold condition.

The model could fit a wide range of knee angle trajectories in the hold condition. The model with personalized parameters predicted the effect of pre-movement demonstrating that the model captured the underlying mechanism and subject-specific deficits.

Our model thus allows us to determine subject-specific origins of joint hyper-resistance and thereby opens perspectives for improved diagnosis and consequently treatment selection in children with spastic CP.

## Introduction

Joint hyper-resistance is a common symptom in neurological disorders (1). Joint hyper-resistance has both neural and non-neural origins. Neural origins are increased muscle tone or background muscle activity and stretch hyper-reflexia. Non-neural origins are changes in mechanical muscle-tendon tissue properties (2). The contribution of neural and non-neural factors varies between patients and knowledge of individual origins of joint hyper-resistance is important to inform treatment selection. However, it has been challenging to distinguish these origins in individual patients based on clinical tests that only provide information on the overall resistance against a passive movement (3,4). Instrumented assessment of joint hyper-resistance in combination with parameter identification based on neuromechanical models might allow us to determine how neural and non-neural origins contribute to joint hyper-resistance in individual patients (5). Yet, such parameter identification requires a neuromusculoskeletal model that accurately represents the underlying mechanism of joint hyperresistance.

Joint hyper-resistance is clinically measured as resistance against a passive muscle stretch. During such tests (e.g., Modified Ashworth or Tardieu Scale), the patient is asked to relax. An examiner rotates the joint under investigation at different speeds and subjectively rates the resistance against the imposed movement (3). Important limitations of such clinical tests are that they depend on the interpretation of the examiner and give little insight in the underlying origins of the observed joint hyper-resistance.

Instrumented spasticity assessments have been introduced with the aim to better distinguish neural and non-neural origins of joint hyper-resistance (5,6). During an instrumented spasticity assessment, joint kinematics, joint torques, and/or muscle electromyography (EMG) are measured while passively rotating the joint (7–11). Joint rotations are either applied manually or by robotic devices. The muscle response to stretch as derived from EMG provides a more direct measure of neural contributions to joint hyper-resistance (8,12,13). When EMG is combined with kinematic (joint angles) and kinetic (joint torques) data, a more comprehensive assessment of joint hyper-resistance is achieved (10,14,15). Although instrumented approaches yield more insight in neural and non-neural origins of joint hyperresistance than clinical tests, it remains difficult to disentangle neural and non-neural origins as underlying parameters such as muscle tone, stretch reflex excitability, and muscle stiffness cannot be directly measured. In addition, there is no simple relationship between these underlying parameters and the outcome measures. For example, the EMG response to stretch depends on reflex hyperreflexia and its interaction with muscle tone (16,17).

Combining instrumented tests of spasticity with neuromechanical models might allow the identification of neural and non-neural origins of joint hyper-resistance (5,7,18–22) but requires models that capture the mechanism underlying joint hyper-resistance. Neuromechanical models are mathematical descriptions of the neuro-musculoskeletal system. In such models, subject-specific properties such as muscle tone, stretch reflex excitability, and muscle properties (e.g., stiffness) are represented by model parameters. Model parameters can be estimated by maximizing the fit between simulated and experimental kinematic or kinetic trajectories. This approach has been applied to asses wrist flexors (18,19), ankle plantarflexors (7,20,21), and knee extensors (22). Proposed approaches differ in the neuromechanical model as well as in the input data that is used for parameter estimation.

Notwithstanding large differences in model complexity and the number of parameters used to represent subject-specific properties (from four to over ten (7,18–22)), most models describe contributions from reflex hyper-excitability and altered mechanical properties to hyper-resistance. Notwithstanding consensus about muscle tone being a contributor to joint hyper-resistance (2), only one of the proposed models (21) accounts for muscle tone. Yet, this model did not capture interactions between muscle tone and reflex activity.

Our recent work suggests that muscle tone and its interaction with reflex activity through muscle shortrange stiffness (SRS) shapes the response to a passive muscle stretch (16,23,24). SRS is a sharp increase in force upon stretch of a muscle that has been held isometric. SRS scales with muscle tone and decreases with prior movement (23,24). In addition, SRS interacts with reflex activity through force encoding in the muscle spindles (25). We found that it was crucial to account for SRS (23,24) and its interaction with reflex activity (25) to simulate the decreased first swing excursion during the pendulum test in children with spasticity cerebral palsy (16). The pendulum test is an instrumented test that has been shown to be sensitive for the presence and severity of quadriceps spasticity (26). During the test, the lower leg of a seated and relaxed patient is dropped from the horizontal position and knee kinematics are recorded. Upon release of the lower leg, the leg swings under the influence of gravity and the quadricep muscle is stretched. In typically developing children, the lower leg behaves as a damped pendulum. With increasing levels of spasticity, as measured by the Modified Ashworth Scale, the first swing excursion and the number of oscillations decrease and the leg comes to a rest in a less vertical position (26). With higher levels of muscle tone, as often seen in children with cerebral palsy (1), SRS force upon stretch increases (23). Importantly, SRS depends on the muscle’s movement history and decreases with previous movement (23,27,28), which explains the disproportionally large effect on the first swing excursion. We also tested the role of SRS experimentally (29). As SRS is movement history dependent (23,24,27), we modulated the presence of SRS by moving the leg down and up before releasing it during the pendulum test. Indeed, we found that after moving the leg (i.e., decreasing SRS), the first swing excursion was increased in both children with cerebral palsy and typically developing children, suggesting SRS contributes to joint hyper-resistance. Furthermore, the increase in first swing excursion was larger in children with cerebral palsy compared to typically developing children, suggesting that the interaction between SRS and muscle tone and reflex activity plays an important role in joint hyper-resistance.

Here, we propose to use a neuromechanical model that accounts for SRS and its interaction with muscle tone and reflex activity to identify subject-specific origins of joint hyper-resistance based on knee kinematics measured during the pendulum test. We performed the pendulum test in two conditions, i.e., a hold condition in which we held the leg still before dropping it, and a pre-movement condition in which we moved the leg down and up before dropping it. To evaluate our model, we first tested its ability to fit a wide range of pendulum test kinematics in the hold condition in 5 typically developing children and 15 children with cerebral palsy. We expected parameters describing baseline tone and reflex excitability as well as parameters describing stiffness and damping to be higher and more variable in children with cerebral palsy than in typically developing children. Second, we evaluated whether estimated baseline tone and reflex gains were higher in children that had a clear EMG response to muscle stretch during the pendulum test. Finally, we evaluated whether the model with personalized parameters could predict the effect of pre-movement on the first swing excursion.

## Methods

### Experimental data

This study was approved by the Ethical Committee of UZ Leuven/KU Leuven (s61641). Twenty children (15 CP and 5 TD) participated in this study (table 1). A legal representative of the participant signed the informed consent and participants older than 12 years signed informed assent in accordance with the Declaration of Helsinki. Children with cerebral palsy were recruited through the cerebral palsy reference center. Pendulum test measurements were planned following the clinical gait analysis that was part of their routine clinical care. All patients were diagnosed as having spastic CP confirmed by a neuro-pediatrician. Following inclusion criteria were used: (1) age between 5 and 17 years; (2) Gross Motor Function Classification Scale (GMFCS) level I–III; (3) no orthopedic or neurological surgery in the previous year; and (4) no Botulinum toxin injections at least 6 months before the measurements of this study. Typically developing children were recruited through colleagues and friends.

**Table 1:**
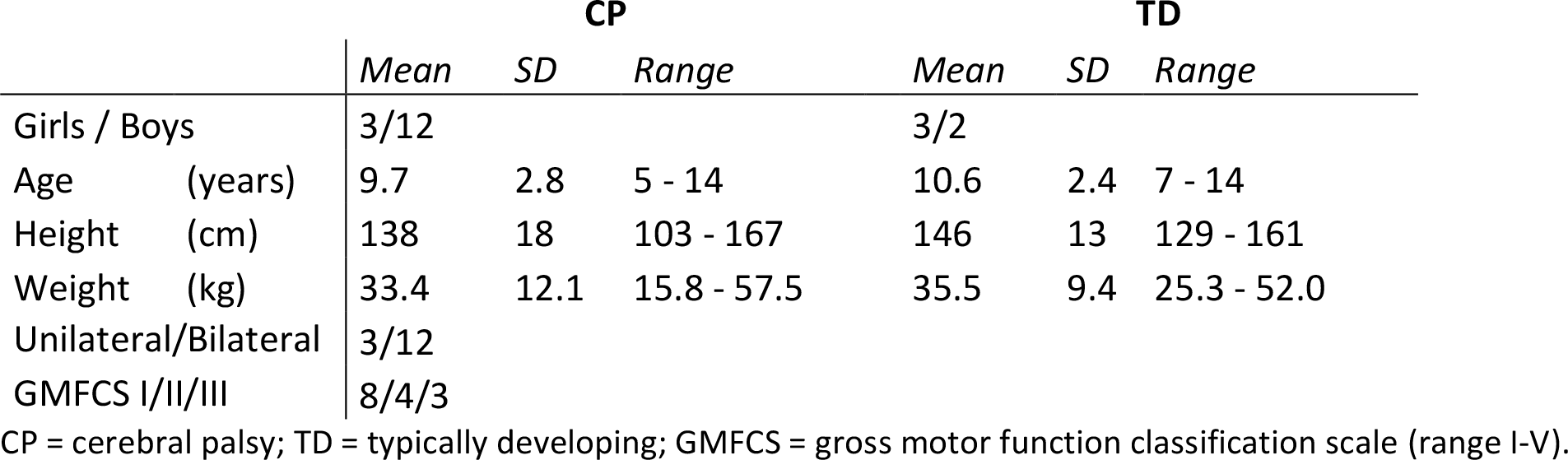
Demographics of participants (mean, standard deviations (SD) and range).

Children with spastic CP were tested in the clinical motion analysis laboratory at the University hospital of Leuven (C-MAL, Belgium). Typically developing children were tested in the Movement and Posture analysis Laboratory Leuven (MALL, Belgium). Both labs are equipped with a Vicon camera system (Vicon, Oxford Metrics, United Kingdom, 100 Hz) for marker-based movement analysis. Reflective markers were placed following the plug-in-gait lower body model (30), extended with markers on the acromion, crista iliaca, trochanter major, and medial knee, malleolus, and toe (see (29) for detailed marker placement). Muscle activity of the rectus femoris, to validate our model, was measured simultaneously by a wireless surface electromyography system (sEMG) (CP: Wave Wireless EMG, Biometrics, United Kingdom, 1000 Hz; TD: ZeroWire EMG Aurion, Cometa, Italy, 1000 Hz) with silverchloride, pre-gelled bipolar electrodes (CP: Nutrode, Xsanatec, Belgium; TD: Ambu Blue Sensor, Ballerup, Denmark). Electrodes were placed according to SENIAM guidelines (31). A custom-made backrest was built to provide back support during the pendulum test (figure 1a).

**Figure 1:**
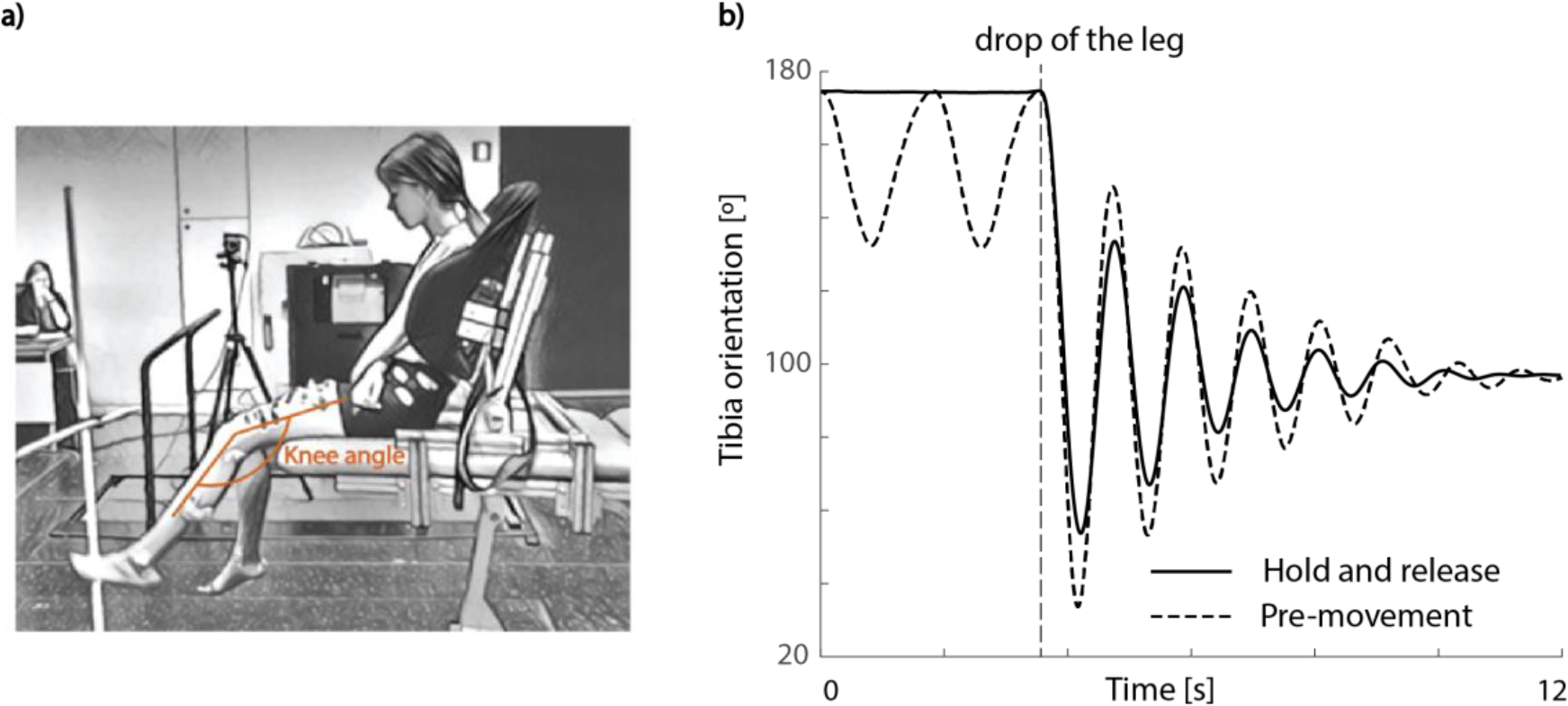
Experimental set-up and exemplar trajectories for the pendulum test. **a)** Children sat with back support from a custom-made backrest during the pendulum test. **b)** Exemplar experimental knee trajectories for the hold and release (full black line) and pre-movement (dotted black line) condition of a typically developing child. The release of the lower leg is indicated with the vertical grey dotted line.

During the pendulum test, the child was asked to sit relaxed with back support (figure 1a). This chair allowed us to control and standardize the hip angle between subjects and trials. On average, the hip was 26° (±11°) into flexion. The examiner extended the leg to a horizontal position and then dropped the lower leg. The pendulum test was performed in two conditions. (1) In the hold and release condition, the leg was held still (isometric) for at least 5s prior to release (figure 1b, full black line). (2) In the movement and release condition (pre-movement) (figure 1b, dotted black line), the leg was moved down and up for at least 5s prior to release.

Marker trajectories from Nexus (Version 2.8.5, Oxford Metrics, United Kingdom) were processed using OpenSim 3.3. A generic musculoskeletal model (Gait 2392) was scaled based on anatomical marker positions using OpenSim’s Scale Tool. Knee joint angles were calculated using OpenSim’s Inverse Kinematics Tool. Finally, tibia orientation in the world was calculated using OpenSim’s Body Kinematics Tool (32,33) and used as experimental input data for parameter estimation. Raw sEMG data were bandpass filtered using a fourth order Butterworth filter between 10 and 450 Hz followed by signal rectification. Finally, a fourth order Butterworth low-pass filter with 20 Hz cut-off was applied.

All trials were inspected for voluntary activity by visually checking muscle activity and kinematic trajectories and trials with signs of voluntary activity were excluded from further analysis (see (29) for detailed exclusion criteria). The most affected leg was evaluated for children with cerebral palsy, while one leg was randomly selected for the typically developing children.

### Neuromechanical model and parameter estimation

We modeled the lower limb and foot as a single rigid body that could rotate at the knee. The knee is actuated by two antagonistic muscle-tendon units, representing the knee extensors and knee flexors. In addition, there was linear joint-level damping. Muscle mechanics were described by a Hill-type model (34,35). For the knee extensor muscle, we extended this model with SRS as this muscle is being stretched following an isometric period (in contrast, the knee flexors first shorten when the leg is dropped during the pendulum test) (figure 2a):

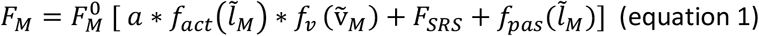

where F is muscle force, 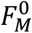 is peak isometric muscle force, *a* is muscle activation,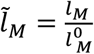 is normalized fiber length, 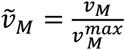 is normalized fiber velocity, and *f*_*act*_, *f*_*pas*_, and *f*_*v*_ are the active muscle force-length, passive muscle force-length, and muscle force velocity characteristics, respectively (35). Muscle short-range stifness, *F*_*SRS*_, was modeled similarly as in De Groote et al (16,36) (equation 2). SRS force was proportional to isometric muscle force at stretch onset and muscle stretch until the critical stretch was reached (i.e., the stretch at which the slope of SRS force sharply decreases in experiments) (24). SRS was constant for the remainder of the stretch.

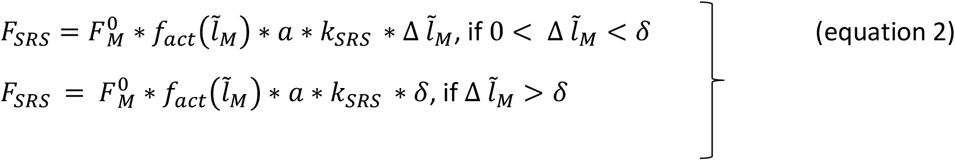

where *k*_*SRS*_ *=* 280, is the SRS constant (16), 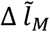 is normalized fiber stretch relative to fiber length at stretch onset, and δ *=* 5.7 ∗ 10^−3^ is the normalized critical stretch (16).

**Figure 2:**
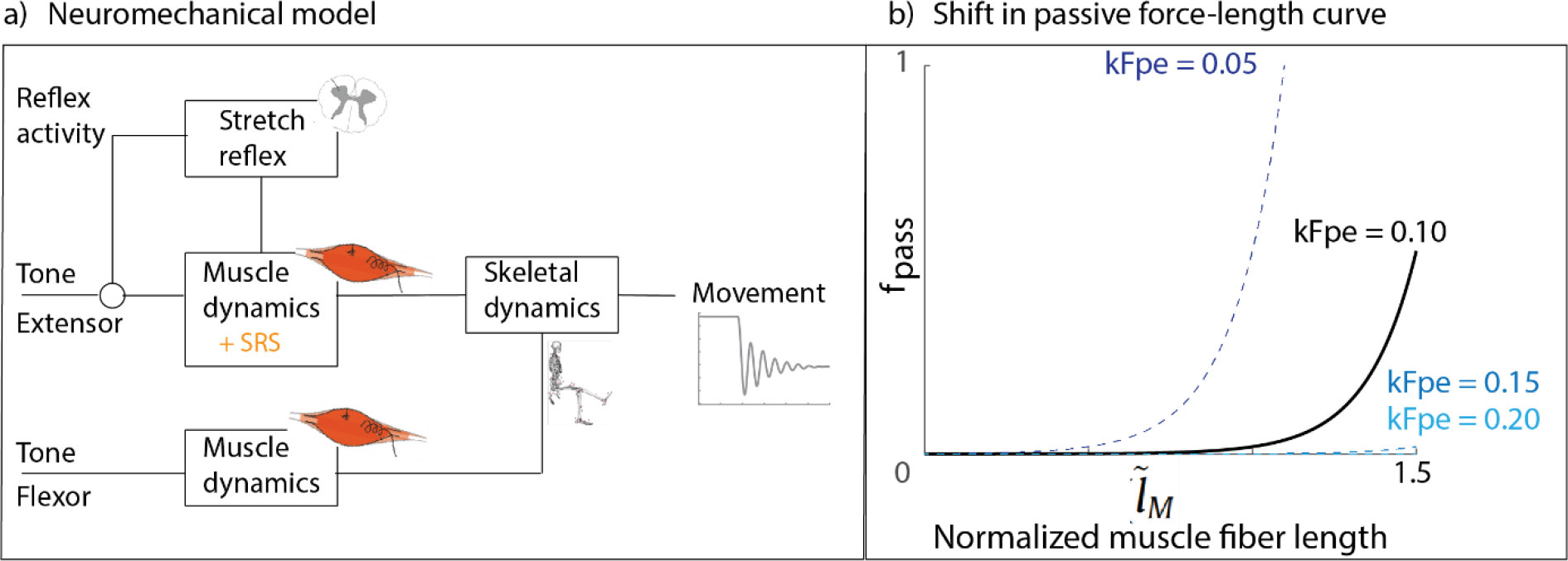
**a)** Schematic representation of our neuromechanical model and **b)** exemplar shifts in the passive forcelength curve. SRS = short-range stiffness; k_Fpe_ is an optimization variable that allows for shifting the curve. Nominal value for k_Fpe_ = 0.10.

We assumed that SRS was only present during the first swing excursion since SRS disappears with prior movement (23,27,28). Furthermore, we assumed that SRS force decayed exponentially when the muscle started to shorten when the leg moved up again after the first swing (equation 3) (16):

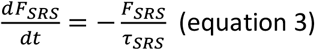

where τ_*SRS*_ *=* 50 *ms*, is the time constant of the exponential decay (16).

Passive muscle stiffness was modeled as an exponential function of muscle fiber length (35) that could be shifted with respect to length (figure 2b).

Musculoskeletal geometry and muscle-tendon parameters were obtained from the rectus femoris (for the extensor) and biceps femoris (for the flexor) of OpenSim’s Gait2392 model scaled to the subject’s anthropometry. Isometric muscle forces were scaled to body weight to the power 0.67 (37). Lower limb inertia was also taken from the OpenSim model, but we allowed up to 20% variation during the optimization to account for differences in inertial parameters between adults and children (38).

Muscle activity consisted of baseline muscle activity and in the case of the knee extensor muscle also of reflex activity (equation 4):

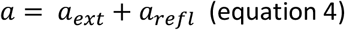

where *a*_*ext*_ is baseline muscle activation for the extensor muscle, and *a*_*refl*_ is reflex activity. Reflex muscle activity was modeled as delayed (τ = 80 ms) linear feedback from active muscle force (including SRS force) (16,25) whenever the muscle force was higher than the muscle force prior to muscle stretch (equation 5):

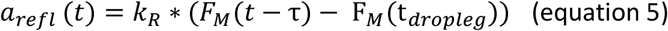

where *k*_*R*_ is the optimized reflex gain, *F*_*M*_ the active muscle force, and *F*_*M*_(*t*_*dropleg*_) the initial muscle force prior to muscle stretch.

We assumed that subject-specific differences in neural control and muscle properties could be described by the following model parameters: baseline activation of the extensor (*a*_*ext*_) and flexor (*a*_*flex*_) muscle, reflex gain (*k*_*R*_), shift in passive force-length curve (*k*_*Fpe*_), and a damping coefficient (B) for linear damping at the knee joint. These parameters were estimated by optimizing the fit between experimental and simulated knee joint angles, *θ*_*exp*_ and *θ*_*sim*_, and angular velocities, 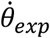 and 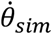. In addition, we penalized high values of baseline muscle activity and reflex gains with a small weight to discourage neural contributions when not needed to fit the measured trajectories (equation 6):

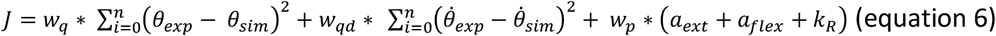

where *w*_*q*_ *=* 1 and *w*_*qd*_ *=* 0.5, are the weights for the error between simulated and experimental angles and angular velocities, respectively; and *w*_*p*_ *=* 0.001, the weight to discourage neural contributions when this was not needed to fit the measured trajectories.

We estimated parameters for each trial of the hold and release condition. The initial state of the knee joint angle and angular velocity were constraint to the experimental initial knee joint angle and zero velocity.

The optimization problems were solved via direct collocation with a trapezoidal integration scheme in CasADi (39), using the interior point solver (IPOPT, solver mumps) (40) with an error tolerance of 10^-7^. To deal with the different dynamics on the first versus consecutive swings, we formulated a two-phase optimization problem. We used N_1_ mesh intervals (with N_1_ the experimental frame number of the end of the first swing excursion) of equal length to describe the state trajectories during the first swing and N-N_1_ mesh intervals of equal length to describe the remainder of the state trajectories with N_1_ being the number of mesh intervals until the end of the experimental first swing excursion and N the total number of mesh intervals. Because the duration of the first swing was unknown prior to solving the optimization problem, the length of the mesh intervals during the first swing was an optimization variable with the constraint that the duration of the first swing was at least 200ms, while the mesh intervals for the remainder of the movement were 20ms. Given that the resulting non-linear optimization problems had many local optima, we solved each optimization problem using ten initial guesses (Supplementary material S1, table S1) and selected the solution that resulted in the lowest cost function.

### Outcome measures

We calculated the root mean square error (RMSE) between the experimental and simulated pendulum test trajectories to quantify how well our model could fit the data.

To validate our model, we evaluated our estimations of neural contributions, i.e. estimated reflex gain and baseline extensor activity, against experimental reflex activity derived from EMG. Note that the EMG data was not used for parameter estimation. First, we divided the experimental trials in a group with no or very low muscle responses and with high muscle responses (low vs. high responsive EMG group) based on the peak (processed) EMG observed during the first swing excursion (when the largest reflex is expected). We used a threshold of 0.01 based on visual exploration (for details, see supplementary material S2, figure S1). We then evaluated whether the estimated reflex gains differed between the low and high responsive EMG group. We repeated this analysis for the reflex gains multiplied by baseline extensor activity as reflex activity interacts with baseline muscle activation. Second, we calculated the Spearman correlation coefficient between experimental peak EMG and estimated reflex gains and between experimental peak EMG and estimated reflex gains multiplied by baseline extensor activity. We evaluated both methods as correlations are sensitive to EMG amplitude and EMG amplitude is sensitive to the experimental conditions. We could not normalize our EMG data as we did not collect EMG during maximum voluntary contractions.

Finally, we compared the experimentally observed and predicted increase in first swing excursion due to pre-movement. Given the variability between trials of the same subject in both the hold and premovement conditions, we calculated the experimental increase in the first swing excursion with premovement between every trial of the hold condition and every trial in the pre-movement condition (e.g., if we had three hold trials and four pre-movement trials, the increase was calculated 12 times (3 HR * 4 MR)). Based on these values, we determined the range and average increase in first swing excursion for every participant. To simulate the effect of pre-movement, we performed forward simulations based on the neuromechanical model from which we removed the SRS force based on the estimated parameters for each hold trial. We then computed the difference in first swing excursion between the corresponding simulated hold and release trials.

Spearman correlations were performed using Matlab (Matlab R2018b, Mathworks, United States) with differences considered significant at p < 0.05.

## Results

### Model performance

We were able to fit a wide range of pendulum test trajectories (RMSE: 3.6° (± 2.1°)) (figure 3a), except for the trajectories of one child (CP 14). Trajectories varied largely across (figure 3b) but also within subjects (supplementary material S3, figure S2).

**Figure 3:**
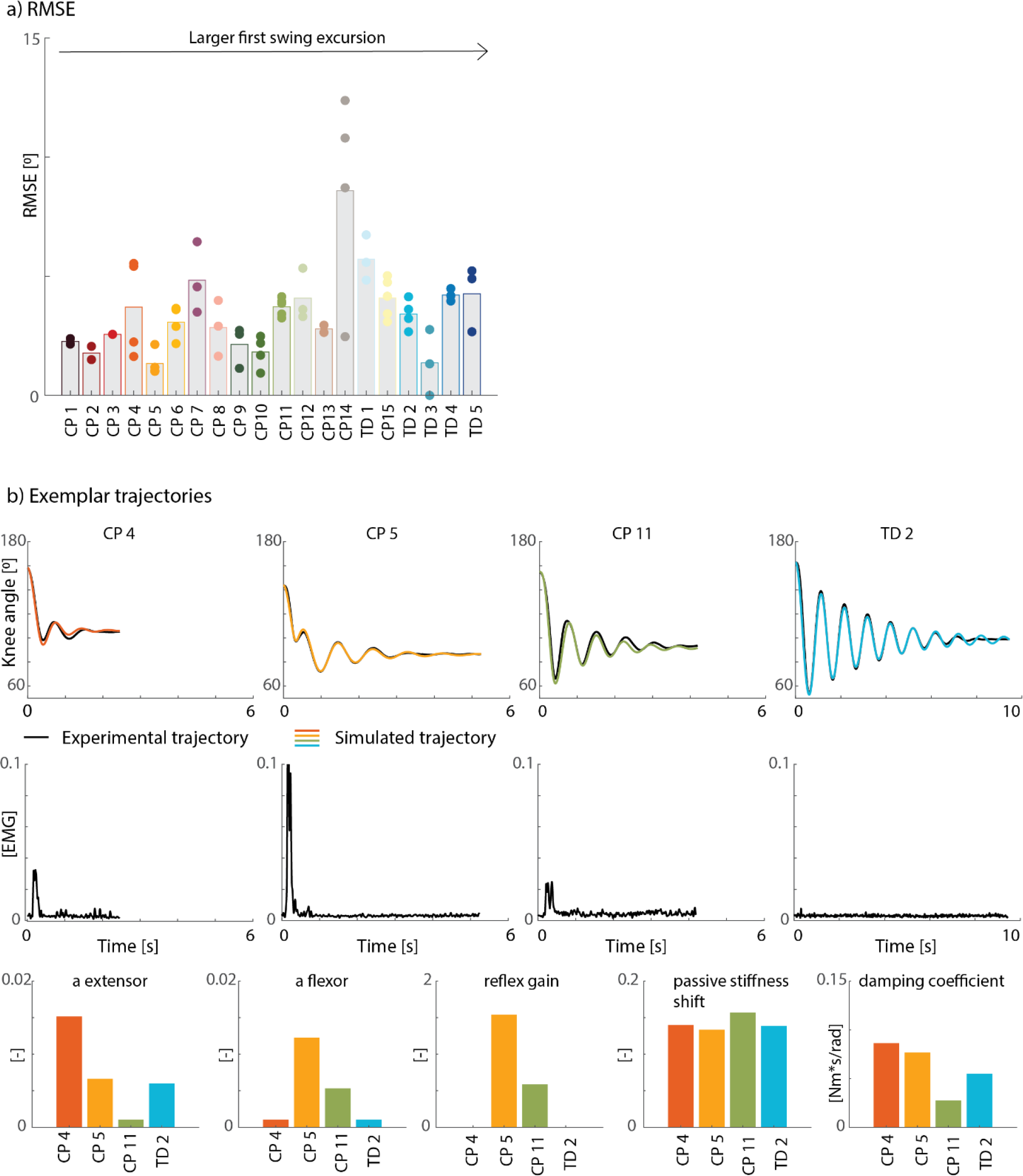
Exemplar experimental data and model fits. **a)** Root mean square errors (RMSE) between experimental and simulated knee ankle trajectories for all children. Children were ranked from low to high experimental first swing excursions. Average values are indicated in bars, RMSE for individual trials are indicated with dots. Every color represents one child. **b)** Exemplar experimental data and simulations for four children (3 CP, 1 TD). Experimental (black) and simulated (color) knee angle trajectories (top row); experimental EMG trajectories for the rectus femoris for the same children and trials (middle row); estimated parameters (bottom row). CP = cerebral palsy, TD = typically developing.

Parameters varied largely across subjects (figure 4) with markedly higher damping, reflex gains, and/or muscle tone for children with a reduced first swing excursion. However, there is no monotonic increase or decrease of any parameter with increasing first swing excursion, suggesting that the origins of a reduced first swing excursion differ between individuals.

**Figure 4:**
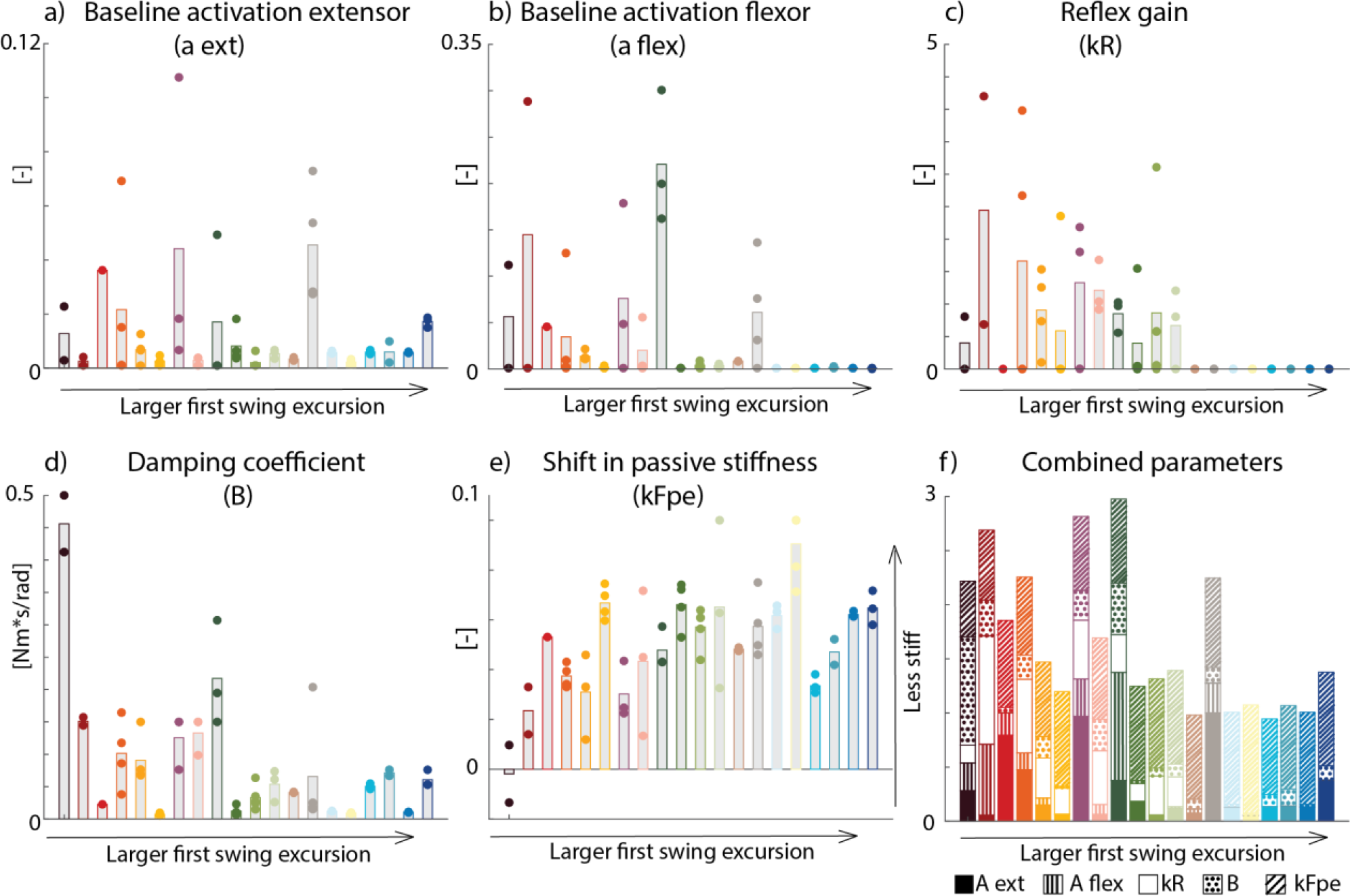
Estimated parameters for all children and all trials (panel a-e). Average values are indicated by the bars, parameters for individual trials are indicated with dots. We subtracted the nominal value of 0.10 from the shift in passive stiffness. Note that higher values indicate lower stiffness. Combined estimated parameters (panel f) indicate different origins contributing to the reduced first swing excursion. Each parameter value is normalized to the maximum value observed across all participants for that specific parameter. Children are ranked following increasing experimental first swing excursion. Typically developing children are represented in tones of blue (right on the x-axis).

### Model validation

Due to technical problems with the EMG system, we had to exclude the EMG data of 2 subjects (no signal was recorded for CP 7 & TD 1).

Estimated neural contributions reflected the presence of responsive muscle activity during the first swing. Estimated reflex gains and estimated reflex gains times baseline extensor activation were low in the group with low experimental peak EMG during the first swing (figure 5a). For the group with high experimental peak EMG activity, the model almost always (10 out of 11 cases) predicted reflex gains. Furthermore, experimental peak EMG was significantly correlated with the estimated reflex gains (r=0.7317; p=0.0006) and estimated reflex gain times baseline extensor activation (r=0.8431, p<0.001) (figure 5b).

**Figure 5:**
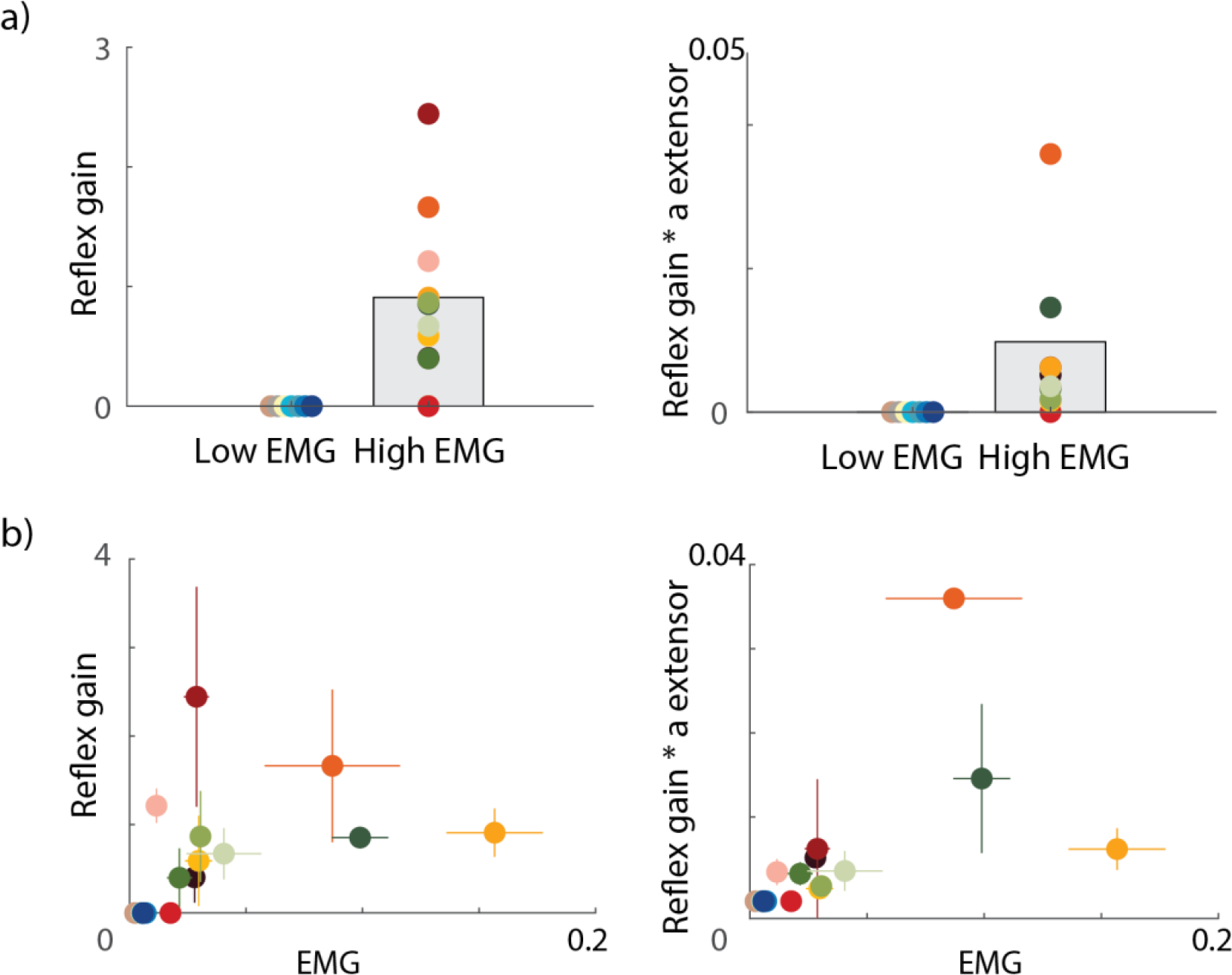
Model validation against experimental EMG data. **a)** Subjects were divided in a low and high EMG response group based on the peak in EMG during the first swing (as a measure for the presence of reflex activity). Estimated reflex gains (left) and reflex gains times extensor baseline activation (right) were plotted for each group. Bars represent average estimated parameter value across all subjects in the low/high group. Every dot represents the average estimated parameter across all simulated trials for one subject. **b)** Correlation between experimental peak EMG and reflex gains (left) and reflex gains times extensor baseline activation (right). Every dot corresponds to one subject with the average peak EMG over all experimental trials on the horizontal axis and the average estimated parameters over all trials on the vertical axis. Standard error for each subject is presented with a colored cross. Typically developing children are represented in tones of blue.

The estimated model parameters have predictive value as they captured the effect of pre-movement on pendulum test kinematics (figure 6-7a). In 43 out of 61 cases (70%), the simulated increase in first swing excursion falls within the experimental range. Furthermore, the average experimental increase in first swing excursion was significantly correlated with the average simulated increase in first swing excursion (r=0.57, p=0.009), (figure 7b).

**Figure 6:**
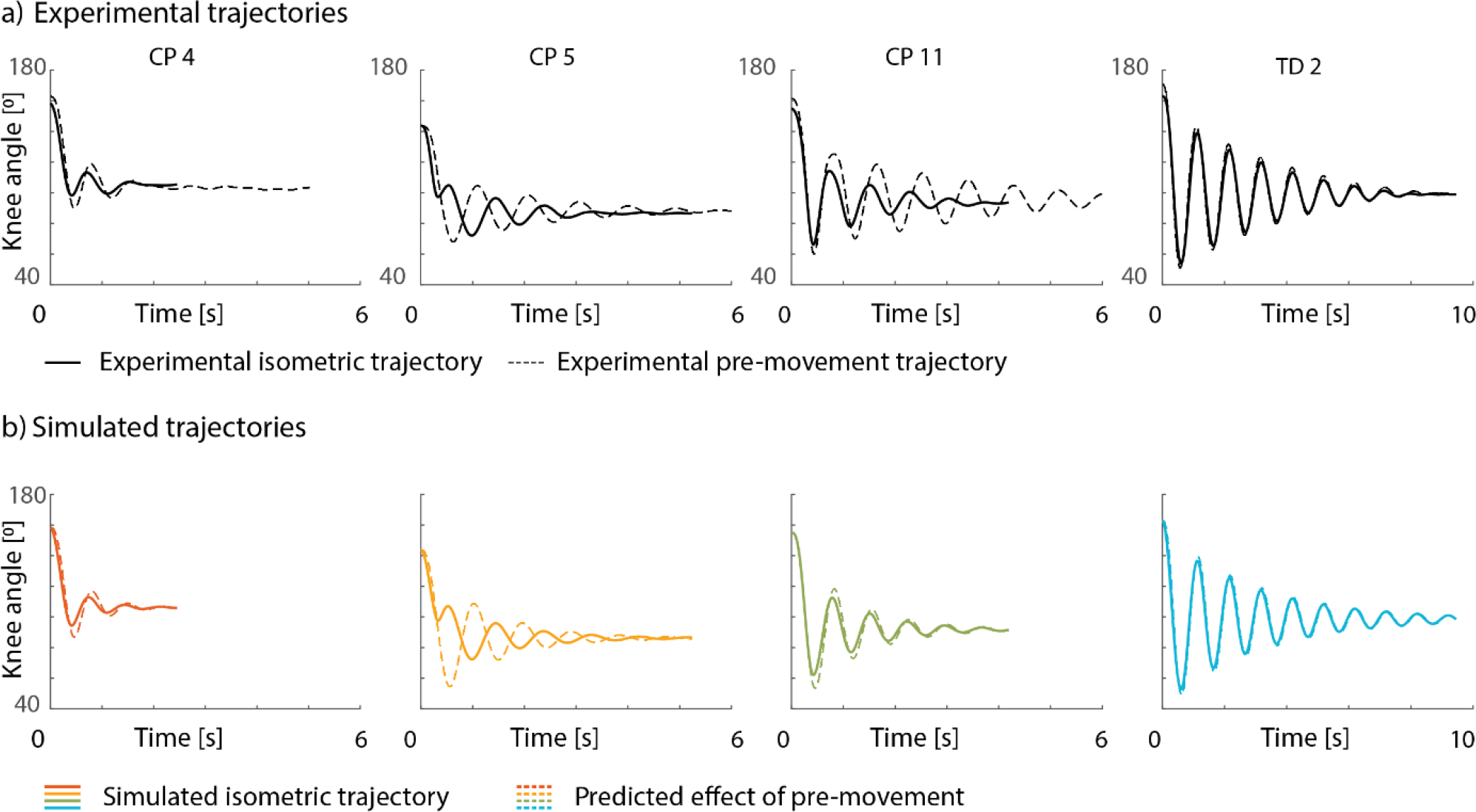
Exemplar trajectories for experimental and predicted effect of pre-movement. **a)** Experimental trajectories. **b)** Predicted trajectories. Isometric trials in full line, pre-movement in dotted line. CP = cerebral palsy; TD = typically developing.

**Figure 7:**
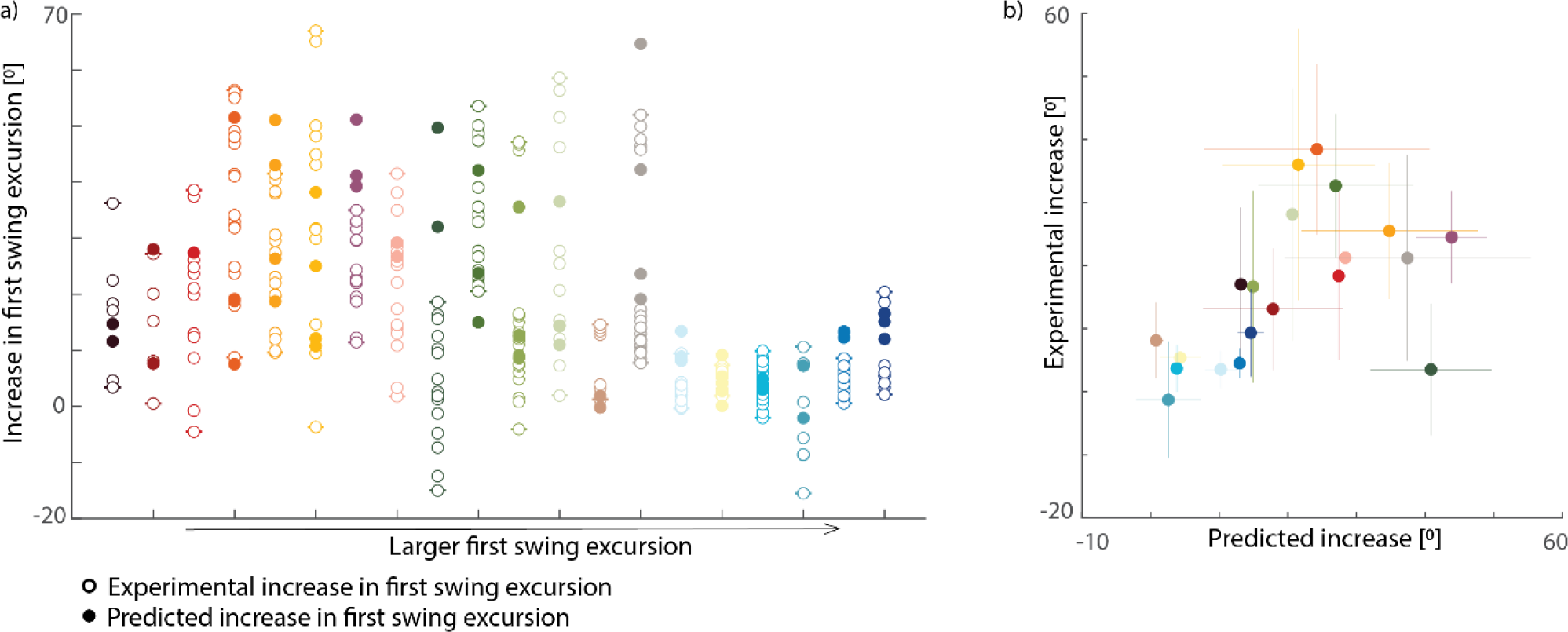
Experimental versus predicted effect of pre-movement on first swing excursion. **a)** Experimental (open dots) and predicted (filled dots) increase in first swing excursion after pre-movement. Subjects are ranked from low to high experimental first swing excursion in the hold and release condition. Note that there are more data points for the experimental condition where each pre-movement trial was compared against each hold trial, whereas in simulation we compared the effect of pre-movement in corresponding trials only (hold and premovement trials based on the same parameters). **b)** Correlation between experimental and predicted increase in first swing excursion after pre-movement. Every dot represents the average (experimental/predicted) increase in first swing excursion across all trials for one subject. Standard error for each subject is presented with a colored cross. Typically developing children are represented in tones of blue.

## Discussion

We can estimate subject-specific neural and non-neural origins of joint hyper-resistance from pendulum test kinematics based on our novel neuromechanical model that accounts for muscle SRS and its interaction with baseline muscle activation and reflex activity. With our model, we could fit a wide range of pendulum test trajectories across typically developing children and children with cerebral palsy with different levels of involvement. The low RMSE values indicate that the measured joint kinematics could be well explained by the model. We found higher neural and non-neural parameters in children with cerebral palsy compared to typically developing children in agreement with known alterations due to cerebral palsy (1). Our approach identified neural contributions for some but not all children in agreement with experimental EMG data, which was not used for parameter estimation. In addition, our model with personalized parameters predicts the effect of movement history, a feature that previous models lacked. Our approach can be applied to other instrumented tests of joint hyper-resistance as long as the joint kinematics and external forces are known. The inclusion of SRS is especially important when tests are performed starting from an isometric condition (27,29), as is the case in many clinical tests of spasticity. We believe that knowledge of the origins of joint hyper-resistance will be useful to elucidate how underlying deficits affect treatment outcome and therefore eventually to improve clinical decision making.

### Model performance and validation

Our model captures mechanisms of joint hyper-resistance that were not described by previous models. Movement history dependent muscle SRS and its interaction with baseline tone have been shown to shape the muscle force response to stretch (16,23,24,27,28). In addition, muscle force seems to drive spindle firing (25). The contribution of SRS is especially large when stretching a muscle that was isometric before the stretch as in the pendulum test but also in other tests of joint hyper-resistance. SRS causes large increases in muscle force for small muscle stretches and thus decouples muscle force changes from muscle length changes. Therefore, modeling reflex activity based on delayed feedback from force might be especially important when SRS is high. It is important that neuromechanical models accurately capture the underlying system dynamics for accurate parameter estimation. In contrast to previous models, we estimated reflex activity using only joint kinematics. Previous models simulated reflex activity based on experimental muscle activity (EMG) or used torque trajectories to estimate reflex activity based on length and velocity feedback (19,22,37).

We successfully identified subject-specific neural and non-neural parameters, which are hard to derive from isolated kinematic features such as the first swing excursion. We found higher neural and/or nonneural parameters in children with smaller first swing excursions compared to typically developing children, which is in line with reported cerebral palsy pathology (1). As there is no monotonic increase or decrease of a parameter with increasing first swing excursion (figure 3 and supplementary material S4, figure S3), the reduction in first swing excursion can have multiple origins. Therefore, underlying origins of joint hyper-resistance cannot be identified based on the first swing excursion alone.

We could predict the effect of pre-movement on pendulum test kinematics based on the parameters estimated based on data collected in the hold condition, suggesting that these parameters capture the underlying impairments and are not merely fitting the data. Estimating the effect of movement history dependency might be of clinical importance as joint hyper-resistance is often measured by stretching a relaxed muscle that has been isometric for some time, while movement history differs across different functional movements, and the magnitude of reflexes is dependent on muscle activity (36,37). This might explain why it has been so hard to interpret how joint hyper-resistance affects daily life activities (e.g., walking) (41,42). The effect of pre-movement is expected to be high when muscle tone and thus SRS is high and when reflex excitability is high. In these cases, we expect that joint hyperresistance will be smaller during movement than during clinical tests and therefore knowing the effect of pre-movement might help predicting the effect of an intervention on function.

### Model considerations

Estimated model parameters differed across trials of the same individual, which was expected based on the large inter-trial variability in knee angle kinematics. However, we expected the parameters representing neural processes to vary between trials but not the parameters representing mechanical tissue properties. The child being more or less relaxed or anxious might change neural parameters (43) and therefore pendulum trajectories whereas tissue properties are constant across trials. We evaluated whether we could explain different trials of a single subject with constant non-neural parameters by estimating trial-specific neural parameters while constraining the damping to the average value of the estimated damping across trials of the same subject. We found that using a constant damping value across trials led to a considerable increase in cost and thus a worse fit and considerably different estimations of extensor baseline activity. The increase in cost suggests that estimated baseline activation and damping are not interchangeable. Our interpretation of this results is that the estimated damping on joint level might consist of the actual joint damping combined with a compensation for the muscle model not accurately capturing activation-dependent damping in the muscle. The activation dependent damping of the Hill-type muscle model is based on isotonic and isokinetic experiments. However, muscles react differently when they operate at varying activation levels or velocities. More complex muscle models might be needed to completely represent the underlying muscle mechanics.

Possibly, different model parameters within one subject might also have been a consequence of different locally optimal solutions. Optimization problems underlying the parameter identification were non-linear and we found that there were many local optima. We tried to reduce the effect of local optima by solving the identification problems from 10 different initial guesses (Supplementary material S1, table S1) and selecting the solution with the lowest cost.

Although our model captured the main kinematic features of a wide range of pendulum kinematics, it failed to capture fine details and a few individual responses (supplementary material S3), possibly due to model simplification. First, our model is phenomenological and might therefore not capture fine details of muscle mechanics (44). Using a more mechanistic cross-bridge model that accounts for SRS might improve the accuracy of our model (45–47). Second, we only accounted for one agonistic and antagonistic muscle, while more muscles influence the knee angle trajectories. Furthermore, we only modeled reflexes for the knee extensors whereas knee flexor activity during the pendulum test has been reported, possibly due to impaired reciprocal inhibition (48). Third, we assumed that we could describe muscle properties of children with cerebral palsy by linearly scaling generic muscle properties (except for increased passive muscle stiffness). However, children with cerebral palsy are known to have altered muscle-tendon properties. They have less sarcomeres in series, but with longer sarcomere lengths, shorter fiber lengths, and more compliant and longer tendons (49,50). Finally, we did not take bony deformities that might alter moment arms into account.

### Clinical significance

Different treatments target different underlying origins of joint hyper-resistance (1), yet knowledge of the origins of joint hyper-resistance is typically limited. For example, baclofen (51), Botulinum neurotoxin injections (52,53), and selective dorsal rhizotomy (54,55) aim to reduce neural contributions to joint hyper-resistance, while stretching and casting (56) aim to reduce non-neural contributions. To select the appropriate treatment, accurate identification of underlying parameters is necessary. Future work should investigate whether the response to treatment is related to underlying neural and nonneural parameters.

The neuromechanical model proposed here is not restricted to the pendulum test alone. The pendulum test is a test of spasticity that can be implemented easily in clinical practice. However, the pendulum test measures only joint hyper-resistance of the knee extensor muscles. Robotic devices (8,9) or mechanical manipulators (14) can also be used to elicit different muscle stretches in other muscles and quantify the response to stretch. As long as the muscle is stretched while the patient is relaxed, and interaction forces and joint movement are measured, our model is applicable.

## Conclusion

To conclude, our neuromechanical model combined with an instrumented test of spasticity (e.g., the pendulum test) can successfully identify subject-specific neural and non-neural origins of joint hyperresistance. Importantly, our model with personalized parameters captures experimental observations that were not used for parameter identification such as the muscle’s activity and the effect of premovement, reflecting that the estimated parameters capture underlying impairments rather than merely fitting the data. Our approach may improve diagnosis and guide treatment selection in children with cerebral palsy.

