## Supplementary material for "A neuromechanical model accounting for movement history dependency identifies subject-specific neural and non-neural origins of joint hyper-resistance: A simulation study"

### S1. Initial guess

As our non-linear optimization problems had many local optima, we solved each optimization problem using 10 initial guesses and selected the solution that resulted in the lowest cost function.

Table S1: Different initial guesses that are used to solve the optimization problem.

| | **IG 1** | **IG 2** | **IG 3** | **IG 4** | **IG 5** | **IG 6** | **IG 7** | **IG 8** | **IG 9** | **IG 10** |
|---|---|---|---|---|---|---|---|---|---|---|
| $A_{EXT}$ | 0.0100 | 0.0050 | 0.0010 | 0.0001 | 0.0100 | 0.0500 | 0.0100 | 0.0100 | 0.1000 | 0.0300 |
| $A_{FLEX}$ | 0.0100 | 0.0010 | 0.0005 | 0.0005 | 0.0050 | 0.0100 | 0.0010 | 0.0010 | 0.0500 | 0.0500 |
| $K_R$ | 0.0100 | 0.0010 | 0.0020 | 1.0000 | 1.0000 | 3.0000 | 1.0000 | 5.0000 | 0.1000 | 0.5000 |
| $K_{FPE}$ | 0.1000 | 0.1500 | 0.2000 | 0.2000 | 0.2000 | 0.1500 | 0.1000 | 0.1500 | 0.2000 | 0.2000 |
| B | 0.1000 | 0.0500 | 0.0600 | 0.0300 | 0.0600 | 0.0200 | 0.0500 | 0.0200 | 0.0300 | 0.0300 |
| DT1 | 0.0050 | 0.0100 | 0.0030 | 0.0020 | 0.0050 | 0.0010 | 0.0020 | 0.0020 | 0.0010 | 0.0010 |
| X | Experimental joint kinematics | | | | | | | | | |
| $\dot{X}$ | Experimental angular velocities | | | | | | | | | |

$a_{ext}$ = baseline activation extensor; $a_{flex}$ = baseline activation flexor; kR = reflex gain; kFpe = shift in passive force-length curve; B = damping coefficient; dt 1 = optimized mesh intervals from phase 1; x = state trajectories for joint angle; $\dot{x}$ state trajectories for angular velocities; IG = initial guess.

### S2. Threshold calculation

We divided average peak EMG response into two categories: (1) low EMG response when peak EMG < 0.01, and (2) high EMG response when peak EMG > 0.01. We defined the threshold based on visual inspection of experimental peak EMG data. All typically developing children were grouped in the low EMG response group, suggesting that our threshold makes sense.

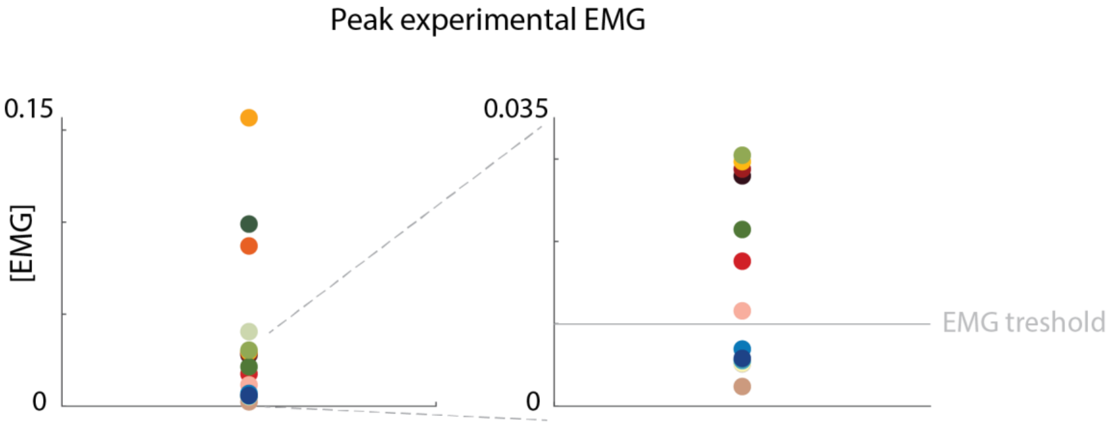


Figure S1: Threshold calculation. Average peak EMG for each subject is represented with one dot (left). Right figure is a close-up from the right figure.

### S3. Pendulum simulations for all participants and all trials

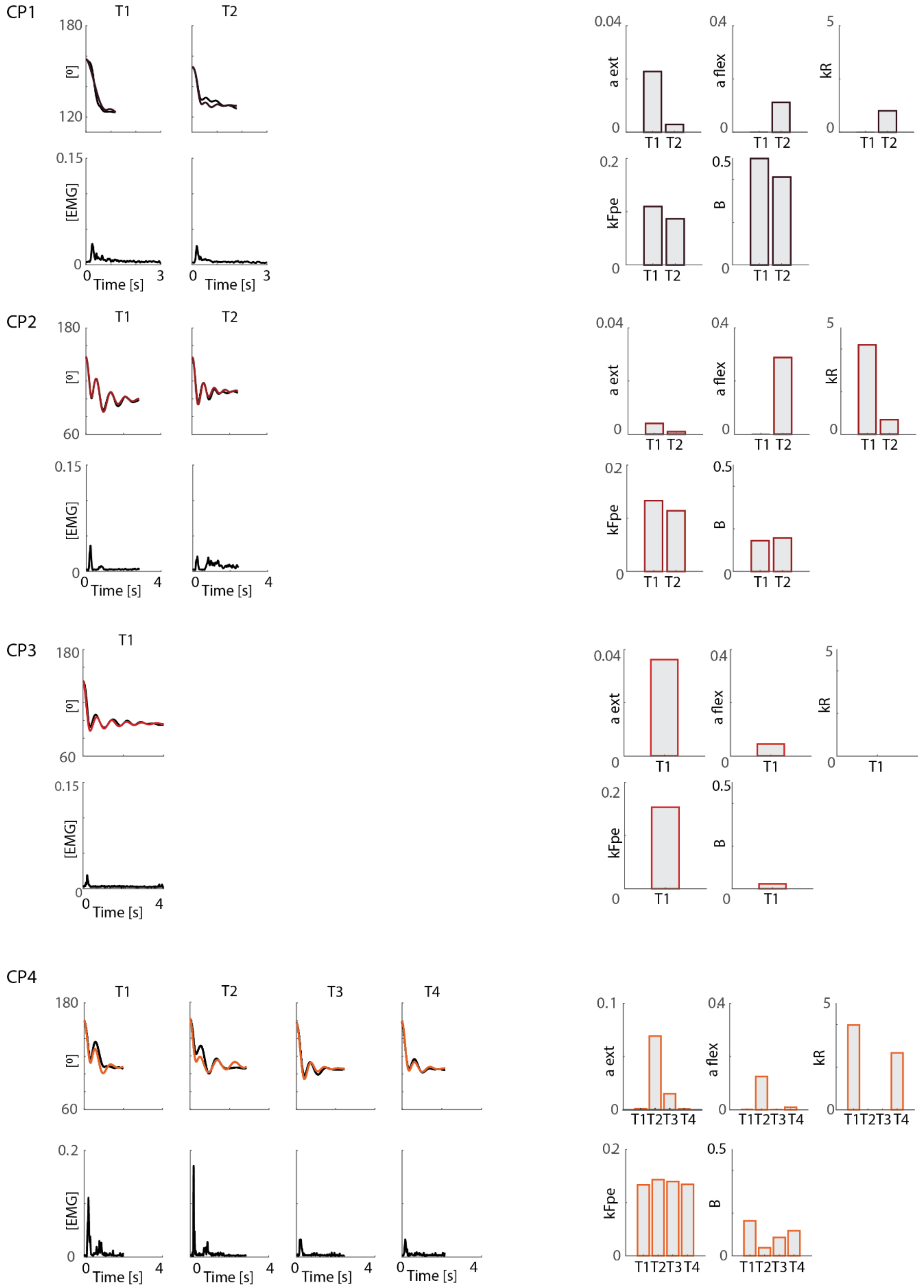


Figure S2: Experimental pendulum and EMG trajectories (black), simulated pendulum trajectories (color) and simulated parameters (right). A ext = baseline muscle tone for the extensor; a flex = baseline muscle tone for the flexor; kR = reflex gain; kFpe = shift in passive length-force curve; B = damping. (Part 1/5)

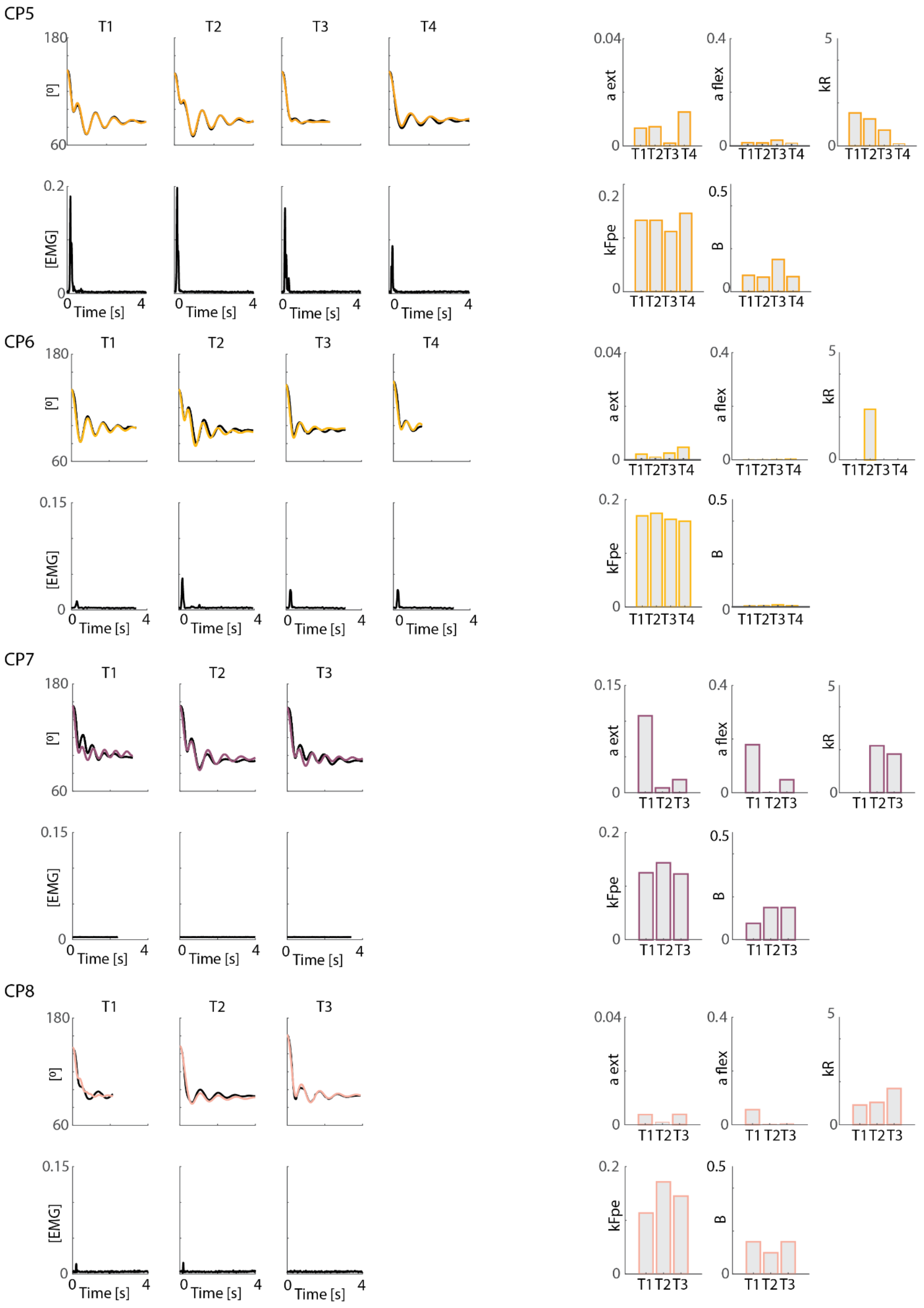


Figure S2: Experimental pendulum and EMG trajectories (black), simulated pendulum trajectories (color) and simulated parameters (right). A ext = baseline muscle tone for the extensor; a flex = baseline muscle tone for the flexor; kR = reflex gain; kFpe = shift in passive length-force curve; B = damping. (Part 2/5)

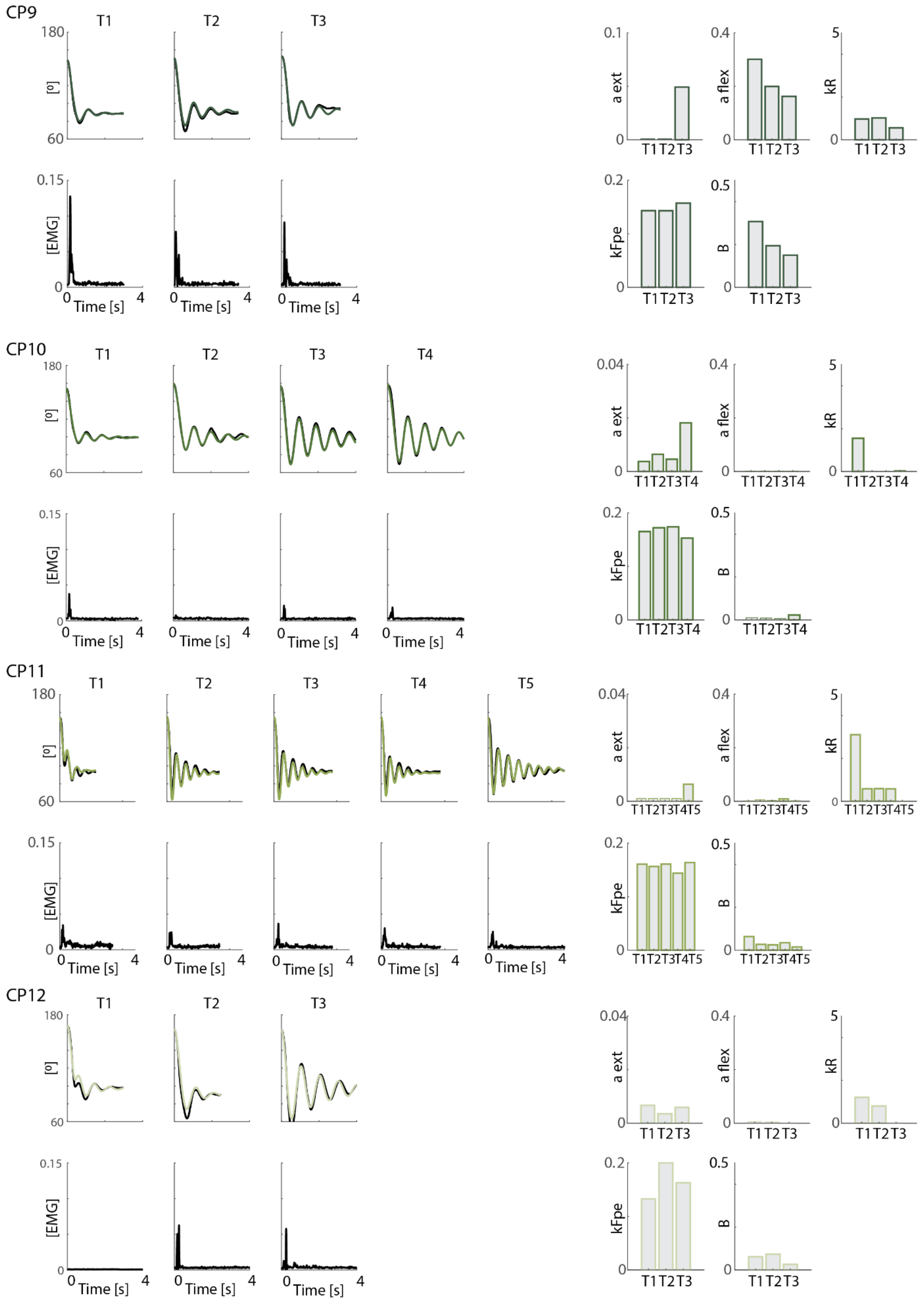


Figure S2: Experimental pendulum and EMG trajectories (black), simulated pendulum trajectories (color) and simulated parameters (right). A ext = baseline muscle tone for the extensor; a flex = baseline muscle tone for the flexor; kR = reflex gain; kFpe = shift in passive length-force curve; B = damping. (Part 3/5)

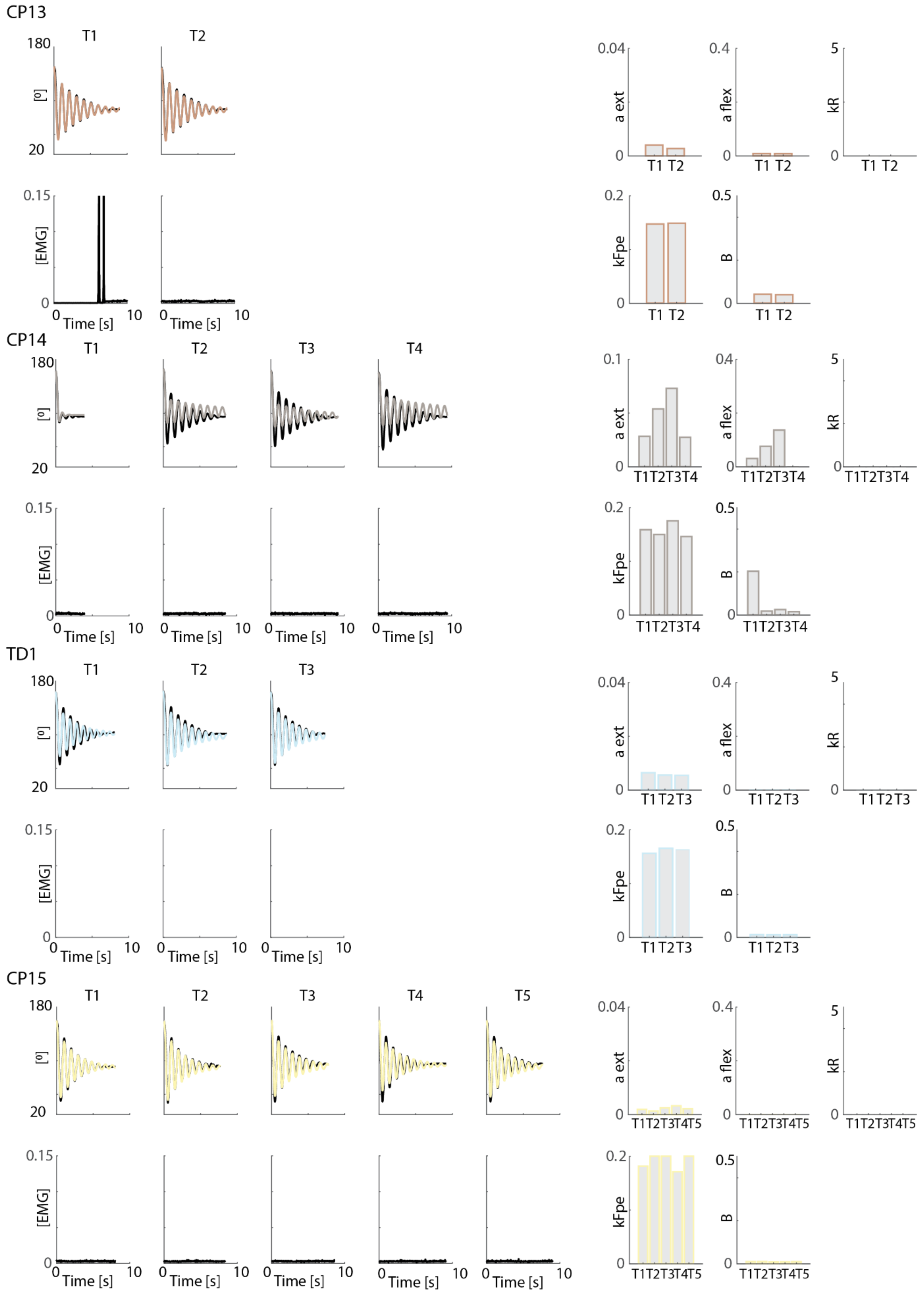


Figure S2: Experimental pendulum and EMG trajectories (black), simulated pendulum trajectories (color) and simulated parameters (right). A ext = baseline muscle tone for the extensor; a flex = baseline muscle tone for the flexor; kR = reflex gain; kFpe = shift in passive length-force curve; B = damping. (Part 4/5)

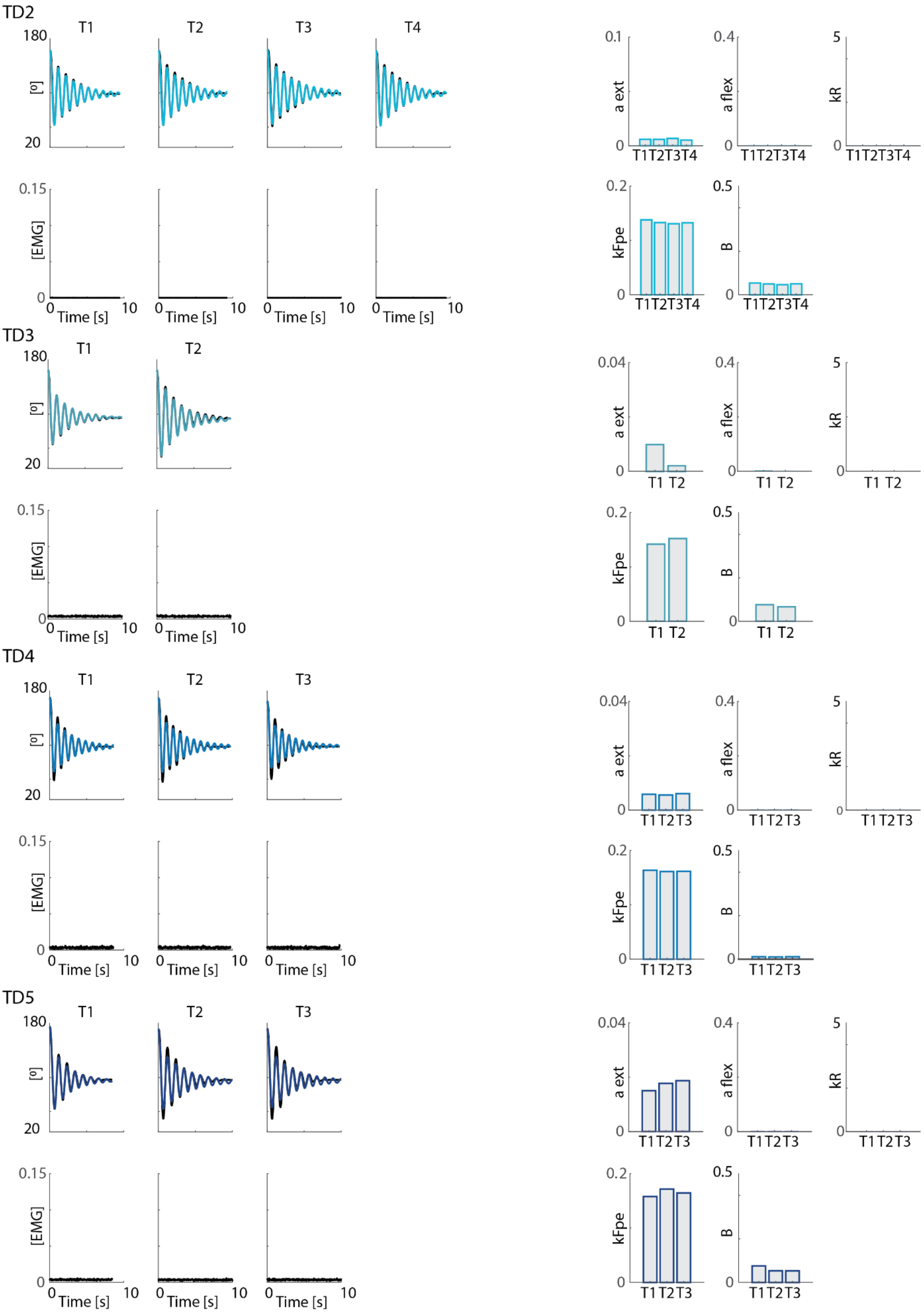


Figure S2: Experimental pendulum and EMG trajectories (black), simulated pendulum trajectories (color) and simulated parameters (right). A ext = baseline muscle tone for the extensor; a flex = baseline muscle tone for the flexor; kR = reflex gain; kFpe = shift in passive length-force curve; B = damping. (Part 5/5)

### S4. Associations between kinematic features and simulated parameters.

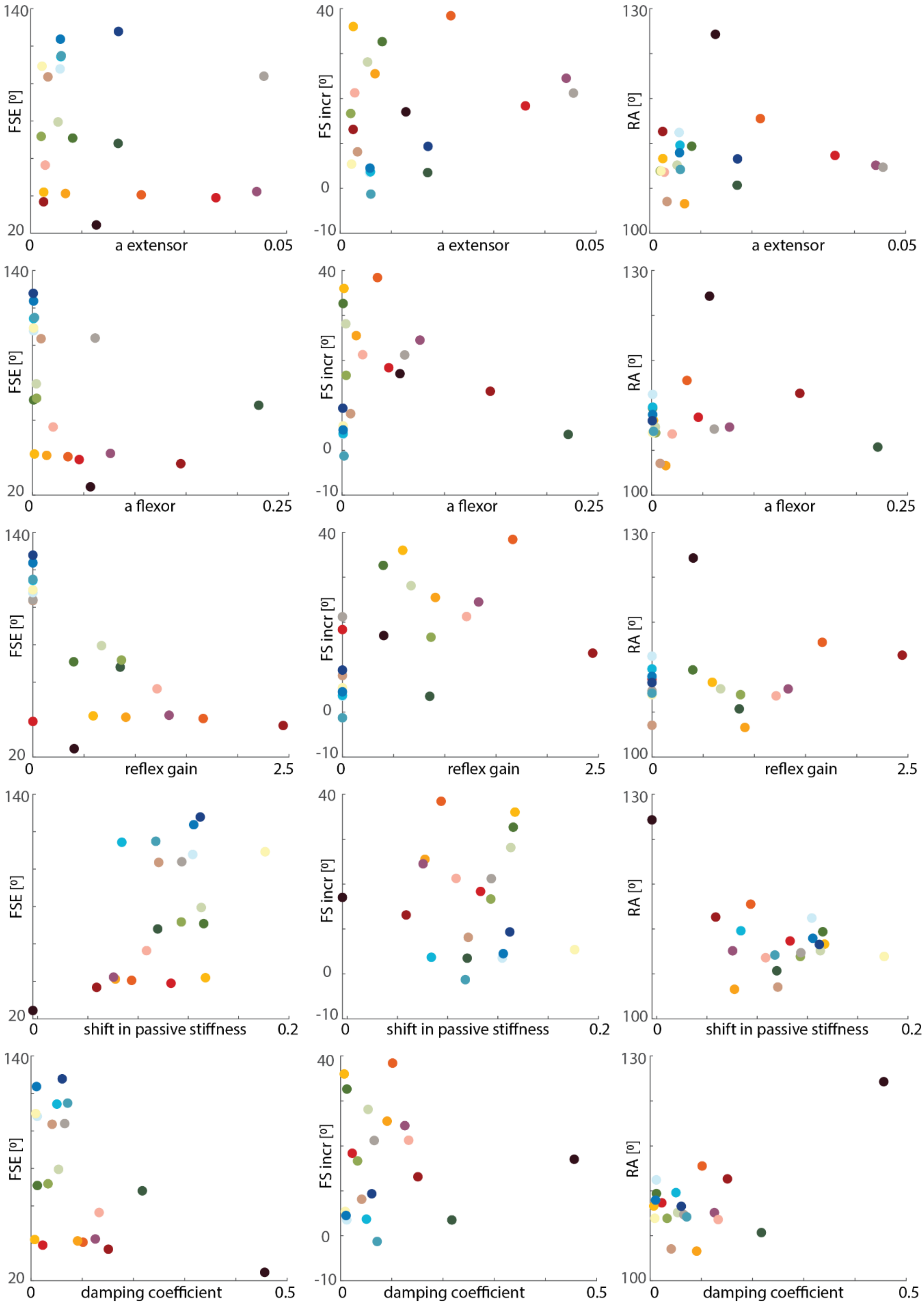


Figure S3: Associations between experimental kinematic features (y-axis) and simulated (x-axis) parameters. Every dot represents the average across all trials for one child. Typically developing children in blue. FSE = first swing excursion; FS incr = increase in first swing excursion after pre-movements; RA = resting angle.

### S5. Post-processing: constant subject-specific damping value.

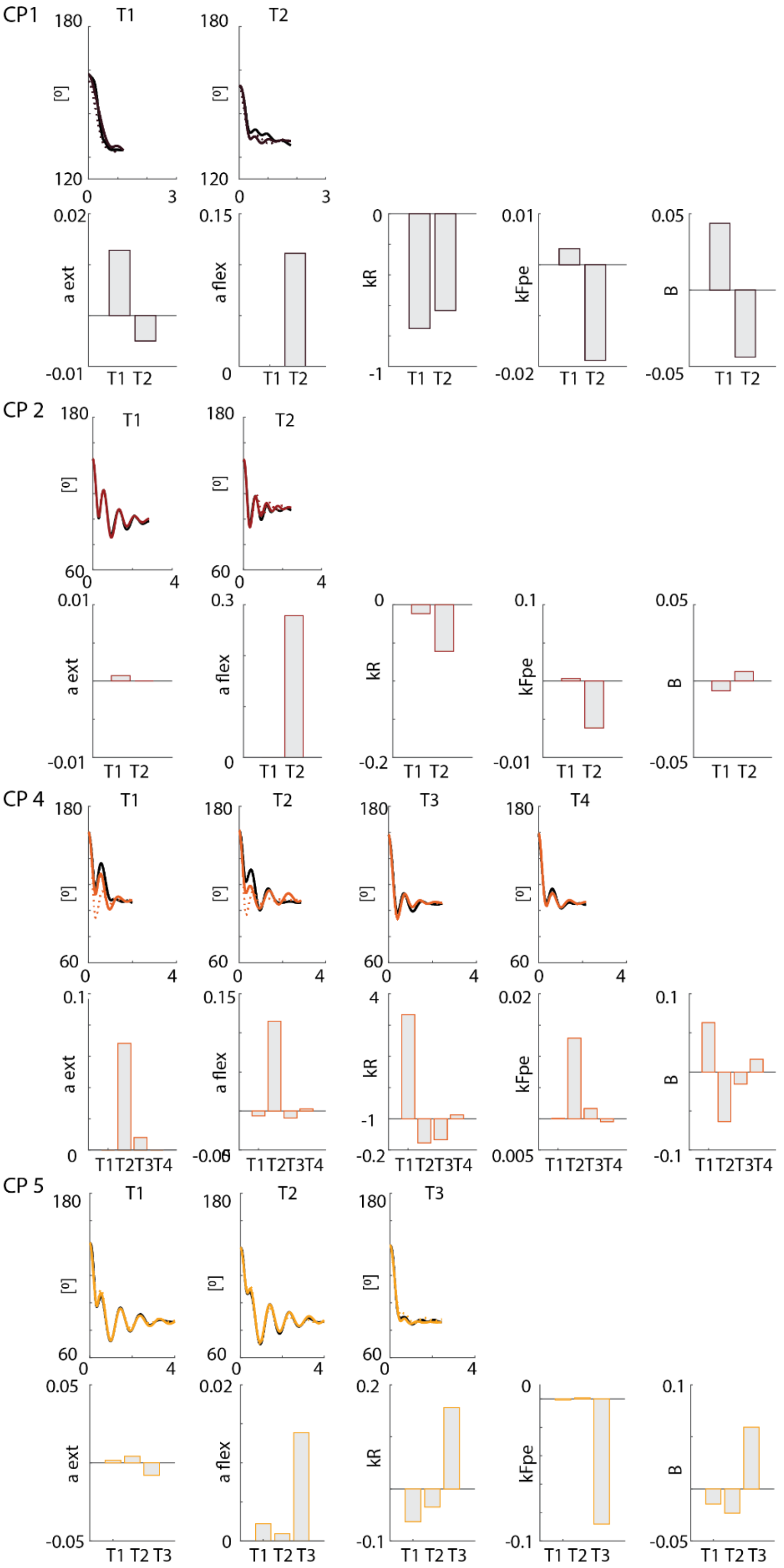


**Figure S4:** Simulated pendulum trajectories with optimized damping (full line) and average damping across trials for one subject (dotted line). Difference between optimized parameters is represented in bars. a ext = baseline muscle tone for the extensor; a flex = baseline muscle tone for the flexor; kR = reflex gain; kFpe = shift in passive length-force curve; B = damping. (Part 1/5)

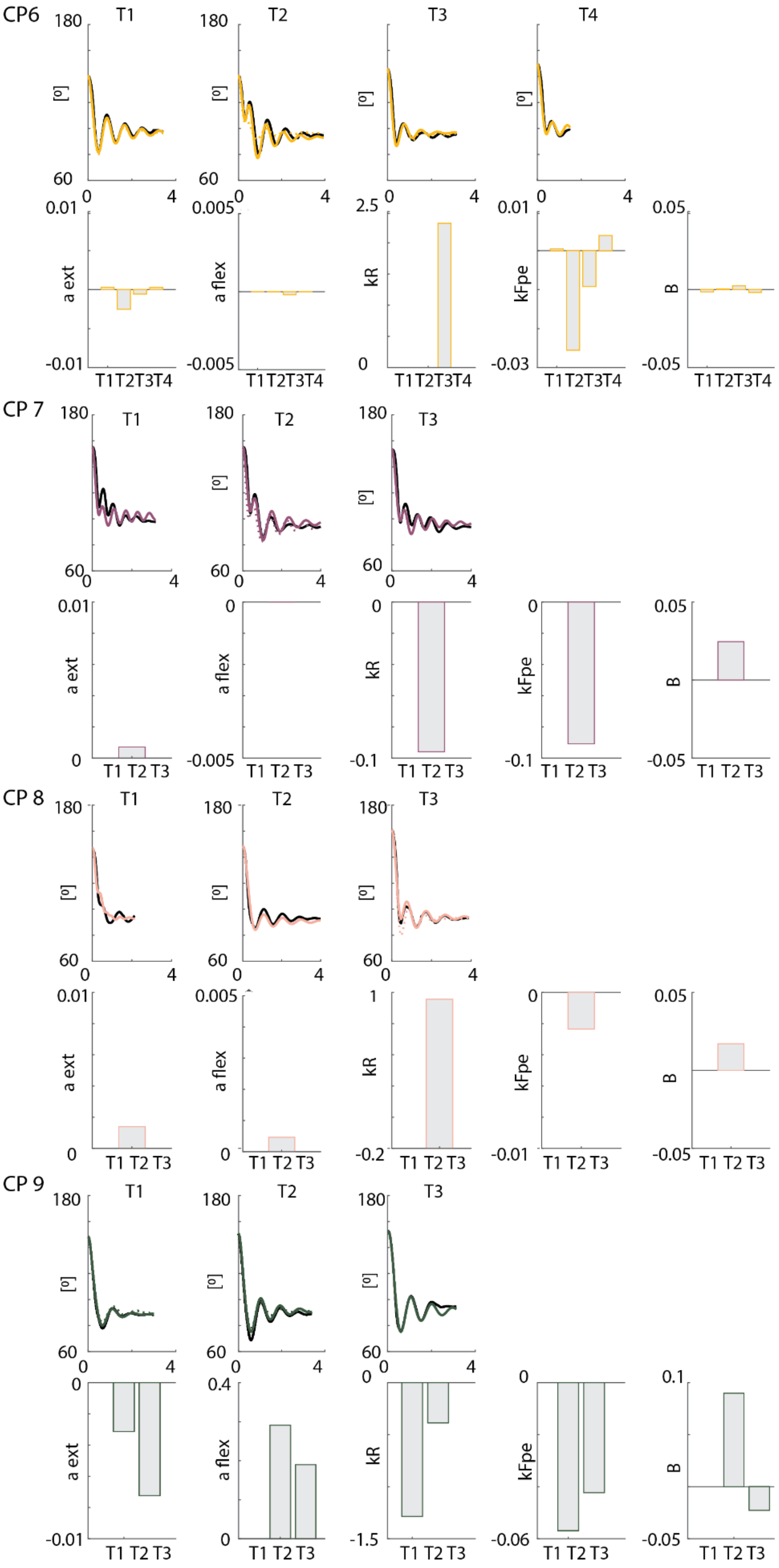


**Figure S4:** Simulated pendulum trajectories with optimized damping (full line) and average damping across trials for one subject (dotted line). Difference between optimized parameters is represented in bars. a ext = baseline muscle tone for the extensor; a flex = baseline muscle tone for the flexor; kR = reflex gain; kFpe = shift in passive length-force curve; B = damping. (Part 2/5)

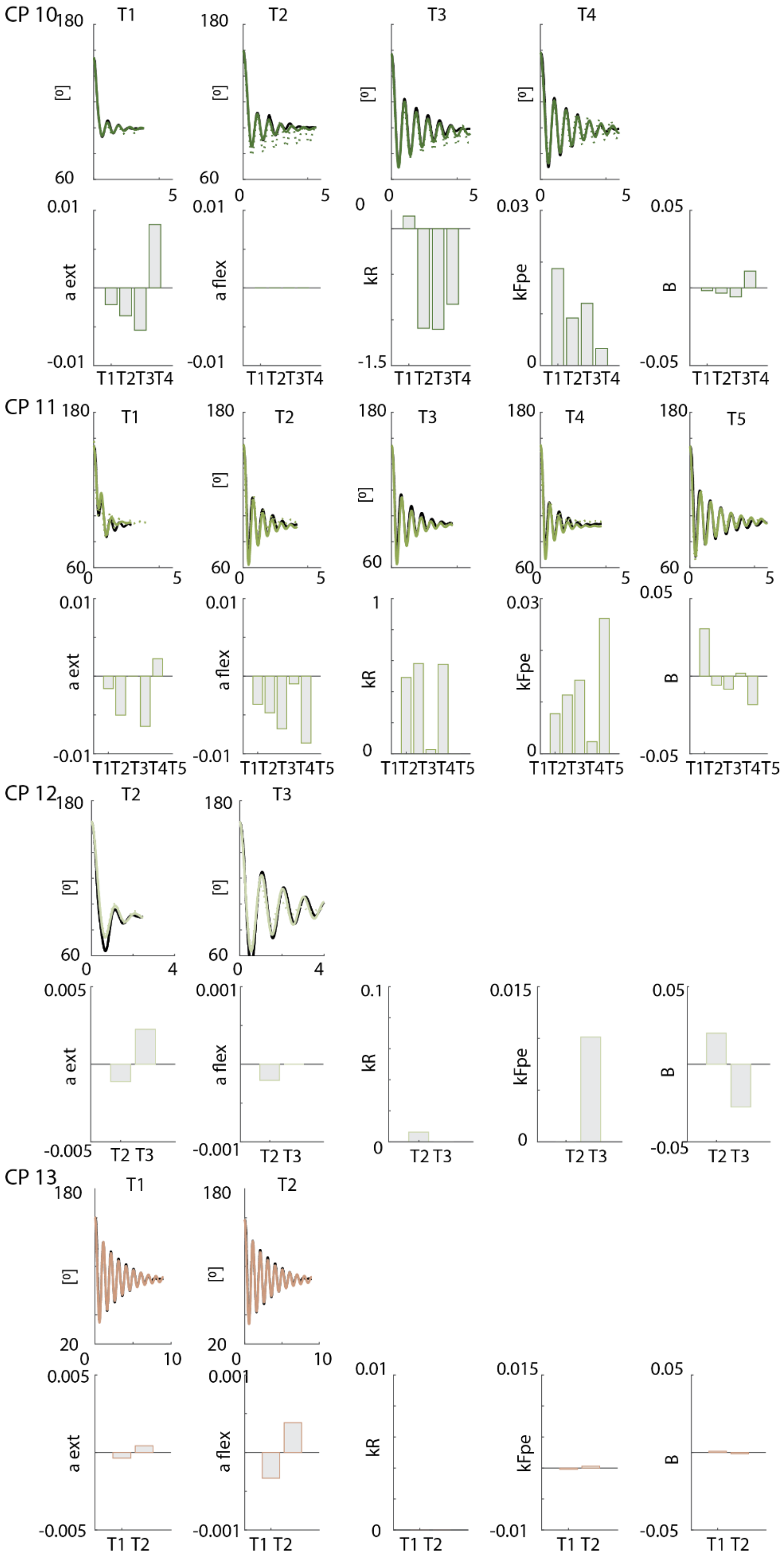


**Figure S4:** Simulated pendulum trajectories with optimized damping (full line) and average damping across trials for one subject (dotted line). Difference between optimized parameters is represented in bars. a ext = baseline muscle tone for the extensor; a flex = baseline muscle tone for the flexor; kR = reflex gain; kFpe = shift in passive length-force curve; B = damping. (Part 3/5)

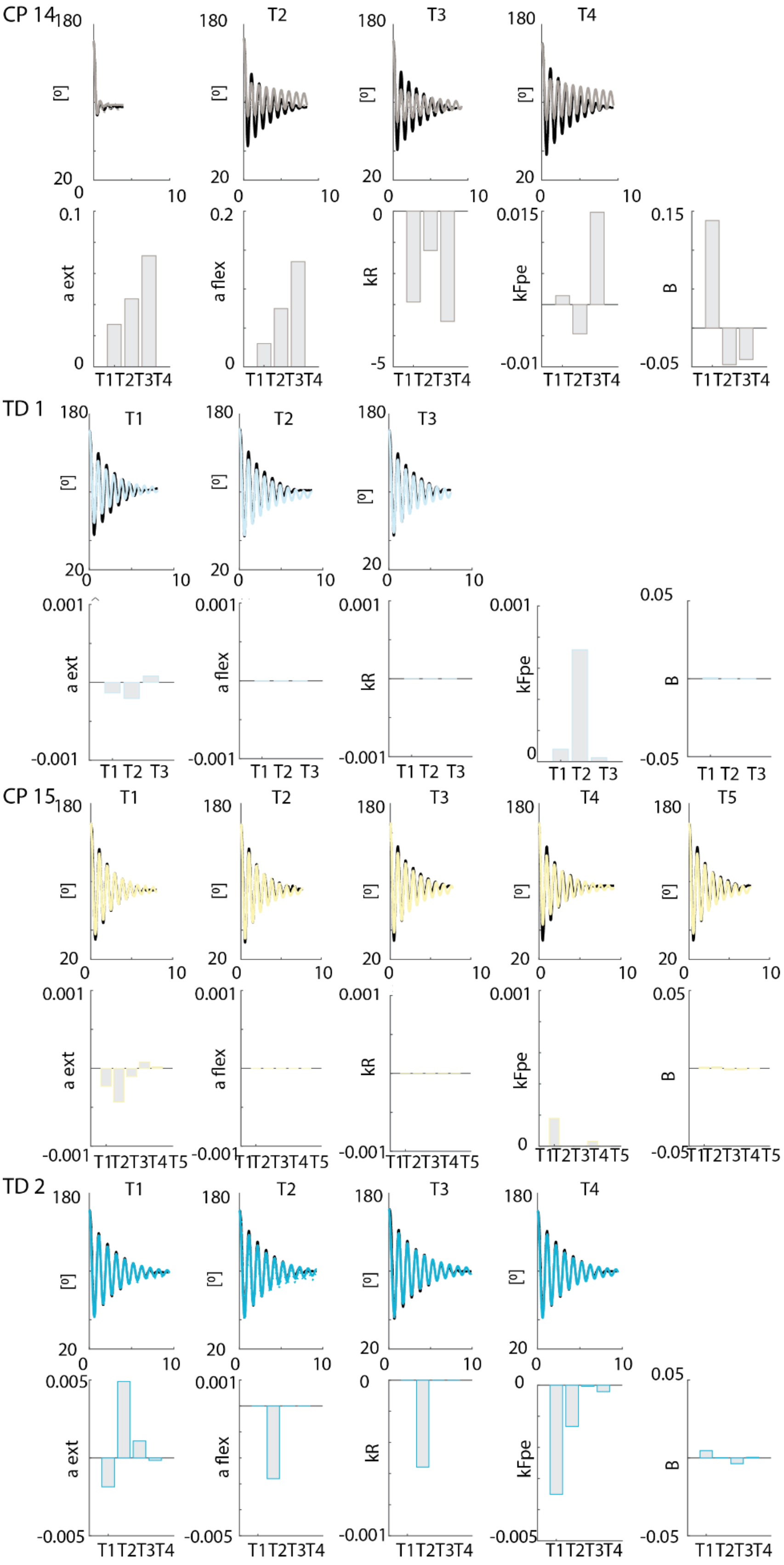


**Figure S4:** Simulated pendulum trajectories with optimized damping (full line) and average damping across trials for one subject (dotted line). Difference between optimized parameters is represented in bars. a ext = baseline muscle tone for the extensor; a flex = baseline muscle tone for the flexor; kR = reflex gain; kFpe = shift in passive length-force curve; B = damping. (Part 4/5)

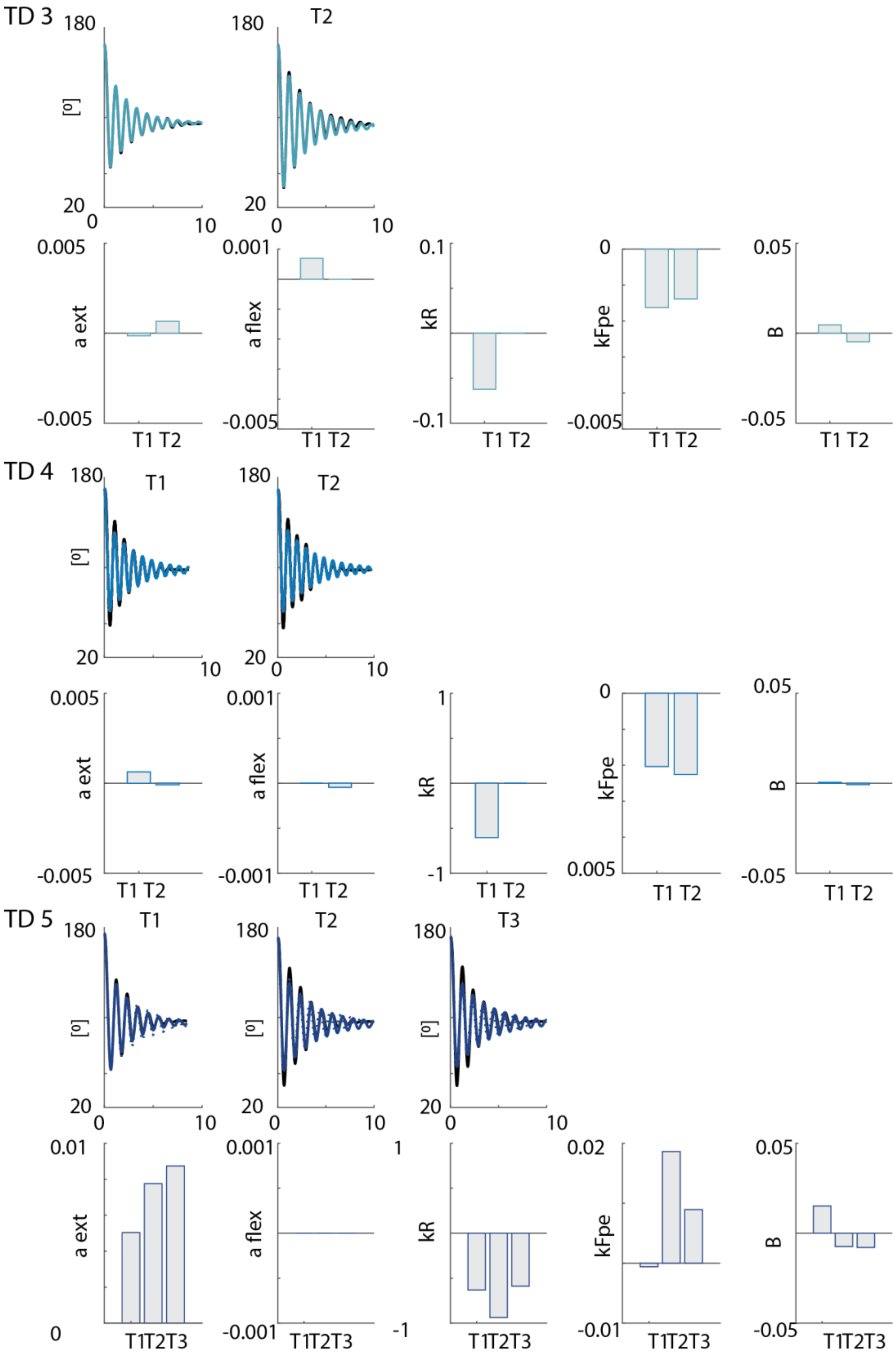


**Figure S4:** Simulated pendulum trajectories with optimized damping (full line) and average damping across trials for one subject (dotted line). Difference between optimized parameters is represented in bars. a ext = baseline muscle tone for the extensor; a flex = baseline muscle tone for the flexor; kR = reflex gain; kFpe = shift in passive length-force curve; B = damping. (Part 5/5)
